# Epigenetic heterogeneity after *de novo* assembly of native full-length Hepatitis B Virus genomes

**DOI:** 10.1101/2020.05.29.122259

**Authors:** Chloe Goldsmith, Damien Cohen, Anaëlle Dubois, Maria-Guadalupe Martinez, Kilian Petitjean, Anne Corlu, Barbara Testoni, Hector Hernandez-Vargas, Isabelle Chemin

**Author notes:** Co-corresponding Authors, **Cancer Research Center of Lyon (CRCL), INSERM U1052, Centre Léon Bérard, 4th floor Cheney B, 28 Rue Laennec, 69373 Lyon Cedex 08, France**.

## Abstract

Hepatitis B Virus (HBV) is a 3.2KB DNA virus that causes acute and chronic hepatitis. HBV infection is a world health problem, with 350 million chronically infected people at increased risk of developing liver disease and hepatocellular carcinoma (HCC). Methylation of HBV DNA in a CpG context (5mCpG) can alter the expression patterns of viral genes related to infection and cellular transformation. Moreover, it may also provide clues to why certain infections are cleared, or persist with or without progression to cancer. The detection of 5mCpG often requires techniques that damage DNA or introduce bias through a myriad of limitations. Therefore, we developed a method for the detection of 5mCpG on the HBV genome that does not rely on bisulfite conversion or PCR. With cas9 guided RNPs to specifically target the HBV genome, we enriched in HBV DNA from Primary Human Hepatocytes (PHH) infected with different HBV genotypes, as well as enriching in HBV from infected patient liver tissue; followed by sequencing with Oxford Nanopore Technologies MinION. Detection of 5mCpG by Nanopore sequencing was benchmarked with Bisulfite-quantitative Methyl Specific PCR (BS-qMSP). 5mCpG levels in HBV determined by BS-qMSP and Nanopore sequencing were highly correlated. Our Nanopore sequencing approach achieved a coverage of ∼2000x of HBV depending on infection efficacy, sufficient coverage to perform a *de novo* assembly and detect small fluctuations in HBV methylation, providing the first *de novo* assembly of native HBV DNA, as well as the first landscape of 5mCpG from native HBV sequences. Moreover, by capturing entire HBV genomes, we explored the epigenetic heterogeneity of HBV in infected patients and identified 4 epigenetically distinct clusters based on methylation profiles. This method is a novel approach that enables the enrichment of viral DNA in a mixture of nucleic acid material from different species and will serve as a valuable tool for infectious disease monitoring.

## Introduction

HBV infection is divided into five clinical categories: asymptomatic, acute, chronic, fulminant, and occult. Occult HBV infection has been defined as the “presence of HBV viral DNA in the liver (with or without detectable HBV DNA in serum) of HBsAg-negative individuals tested with the currently available serum assays” (Raimondo et al. 2010, 2019). Although the mechanism is not well understood, occult HBV infection is associated with liver pathogenesis, significantly increasing the risk for liver cirrhosis and hepatocellular carcinoma (HCC) (Ikeda et al. 2009; Mak et al. 2020; Perumal Vivekanandan et al. 2008). In addition, the overall degree of viral replication has been strongly linked to carcinogenesis (Chen 2006). Therefore, understanding factors that regulate HBV replication may provide insights into occult infection and preventing HCC.

HBV has a particular replication cycle that involves protein-primed reverse transcription of an RNA intermediate called pregenomic RNA (pgRNA) occurring in the nucleocapsid (Wang and Seeger 1993). Upon entry into hepatocytes, viral genomic DNA in the nucleocapsid is in the form of 3.2kb partially double stranded DNA, also called relaxed circular DNA (rcDNA) (Summers, O’Connell, and Millman 1975). The rcDNA is then transported to the nucleus and converted into covalently closed circular DNA (cccDNA) by a still ill-defined mechanism. The HBV polymerase, responsible for reverse transcription, is encoded within the viral genome. The HBV polymerase lacks proofreading activity which results in a high amount of variability; several HBV genotypes have been characterized, named A to J, along with nearly 40 sub-genotypes that have varying characteristics and clinical implications. These different HBV genotypes can vary in their DNA sequence by up to 7.5% (Rajoriya et al. 2017). While our understanding of HBV DNA regulation is still limited, there some evidence that cccDNA could be epigenetically regulated (reviewed in (Xia and Guo 2020).

Recent studies have identified epigenetic modifications of HBV DNA, including methylated cytosines (5mC) as a novel mechanism for the control of viral gene expression (Pollicino et al. 2006). However, the current methods to detect modified bases have a number of limitations. Bisulfite modification has been considered the gold standard for the detection of 5mC for over a decade. This technique converts unmodified cytosines into uracil, and then the 5mC levels are deduced by difference (Li and Tollefsbol 2011). This technique leads to extensive DNA damage and introduces bias through incomplete conversion. Moreover, it is not able to distinguish the difference between 5mC and other modified bases that can occur in the same location (eg 5hmC). Thus, whilst this technique has been incredibly useful, there is a demand for the development of more direct measures of 5mC. Moreover, current sequencing methods are not able to easily distinguish between different forms of HBV (rcDNA verses cccDNA) or identify episomal or integrated HBV DNA.

Nanopore sequencing is a unique, scalable technology that enables direct, real-time analysis of long DNA or RNA fragments (Madoui et al. 2015). It works by monitoring changes to an electrical current as nucleic acids are passed through a protein nanopore. The resulting signal is decoded to provide the specific DNA or RNA sequence. Moreover, this technology allows for the simultaneous detection of the nucleotide sequence as well as DNA and RNA base modifications on native templates (Jain et al. 2016); hence, removing introduced bias from sodium bisulfite treatment and PCR amplification. However, a first enrichment step of the target loci or species prior to sequencing is still necessary. Traditionally, this would be done by PCR amplification, however, this would of course lead to a loss of modified bases like 5mC. Thus, development of enrichment techniques that do not degrade DNA or result in a loss of target bases is needed, in order to fully benefit from Nanopores ability to delineate 5mC levels on native DNA. Nanopore is also capable of sequencing long reads, and can potentially capture the whole HBV genome in single reads and thus provide the sequence information for single HBV molecules. Integration of the entire HBV genome has not been reported, thereofre, sequencing HBV with Nanopore has the potential to distinguish between episomal and integrated HBV DNA. Furthermore, sequencing single HBV molecules could also identify the “gap” in rcDNA allowing discrimination between rcDNA and cccDNA forms of HBV.

In this work we developed a translatable method for the enrichment, sequencing, de novo assembly and the detection of modified bases in the HBV genome; determining the methylation landscape of different HBV genotypes *in vitro* as well as in patient liver tissue. We ascertained the HBV 5mCpG levels, for the first time, on directly sequenced native HBV DNA. These methods represent a valuable and highly novel tool for the detection of modified bases on viral genomes.

## Results

### Enrichment

Performing whole genome sequencing of HBV infected cells without any type of enrichment, achieves an extremely low sequencing depth with any sequencing platform. We performed whole genome sequencing of HBV infected PHH with Nanopore’s MinION and only 1x of HBV was obtained (data not shown). This is due to the size of HBV genome compared to the contaminating host (3.2KB verses 3.2GB, HBV=0.000001% of available), thus enrichment of HBV is clearly necessary. We utilized Cas9 guided Ribonucleic proteins (RNPs) to linearize and enrich in HBV DNA from the total DNA extracted from HBV infected Primary Human Hepatocytes (PHH). This method has been described previously to enrich in target loci in the human genome (Gilpatrick et al. 2020), and to our knowledge, the present study is the first evidence that this approach can be used to sequence viral or circular DNA (figure 1A). This method allows HBV genome enrichment without PCR amplification, thus allowing sequencing of native DNA taking full advantage of the ability of Nanopores to detect modified bases. Briefly, starting material consisted of positive and negative controls for HBV methylation, total DNA extracted from PHH infected with different genotypes (GA, GD or GE) or DNA from fine needle liver biopsies of HBV infected patients (figure 1A). Available DNA ends were blocked by dephosphorylation with calf intestinal phosphatase, this step is essential to prevent the nuclear DNA from being available for the ligation of adapters in later steps. We then used 2 sgRNAs in a highly conserved region, to target the HBV genome, one for the positive strand and an additional guide for the negative strand (figure 1A). Use of 2 gRNAs leading the positive and negative strand was critical, since the RNP complex remains attached to the strand where it makes the cut, making the other strand available for the ligation of adapters allowing the sequencing of both the positive and negative strands. After the adapters and motor proteins were ligated to the newly available DNA ends, the libraries were loaded onto the MinION devise and DNA sequencing occurred in a 5’ to 3’ directionality on a MinION R9.4.1 flow cell.

**Figure 1.**
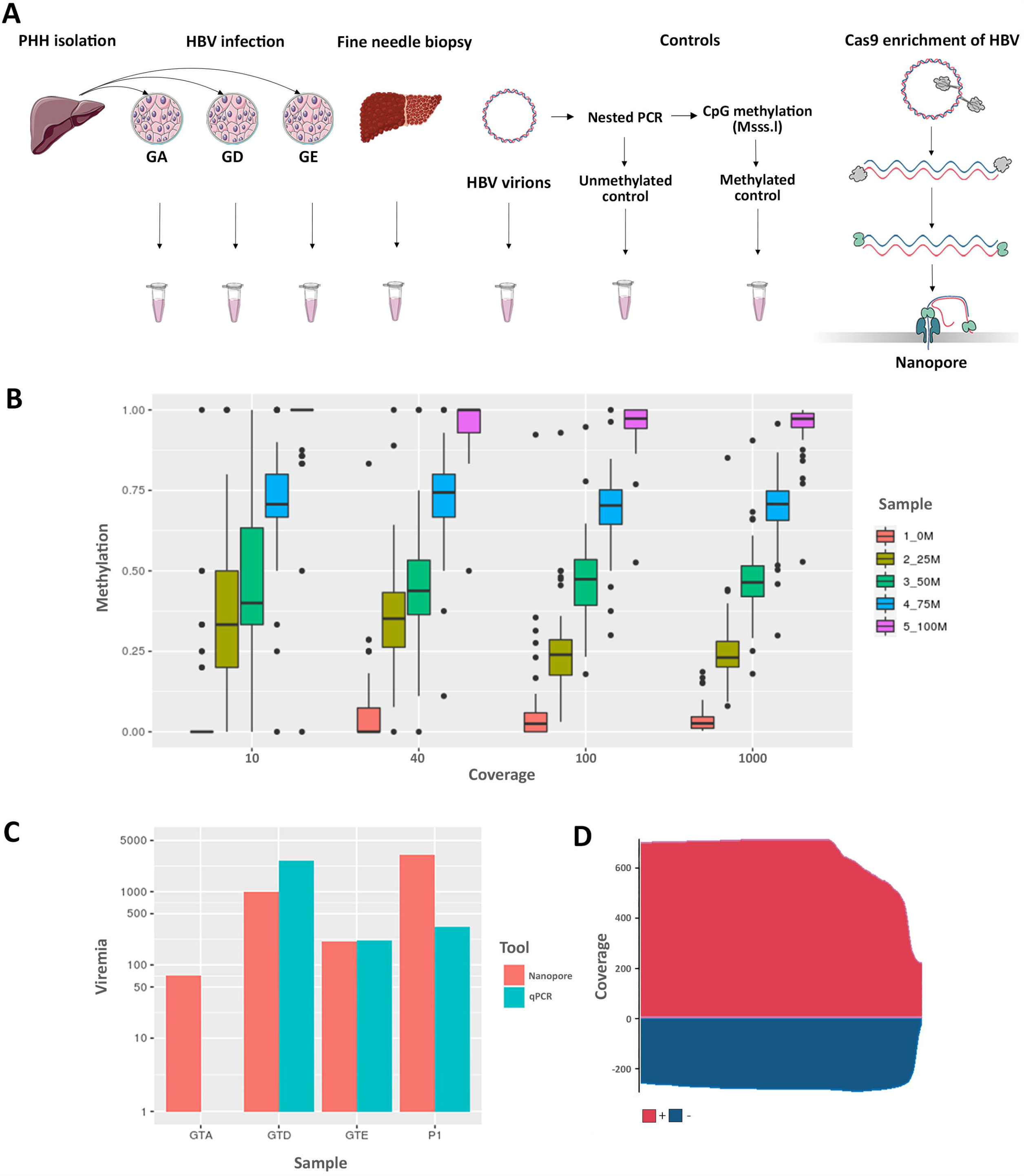
Optimal coverage of HBV for detection of HBV methylation with Nanopore. **A:** Overview of sampling and cas9 targeted sequencing protocol adapted for circular viral genomes. Briefly, all available DNA ends were dephosphorylated prior to liberation of target sites by cutting with cas9 guided RNPs. The circular viral genome was then linearized and prepared for the ligation of adapters and motor proteins. Libraries were then loaded onto a MinION to be sequenced with Nanopores. **B:** calculation of optimal coverage for HBV methylation detection. Briefly, reads from sequencing positive and negative controls for HBV methylation were pooled to achieve 10-1000x coverage and 0, 25, 50, 75 or 100% methylation. Methylation was called with Nanopolish. **C:** Sequencing depth achieved with nanopore (reads aligning to genome) compared to total HBV detected by qPCR (copies of HBV detected by qPCR). **D:** Coverage of HBV genome from patient tissue (P1) after enrichment via cas9 sequencing technique with MinION flow cell, x-axis is HBV genome length (3.2KB).

### Yield and coverage

In order to determine the necessary sequencing depth to detect differential HBV DNA methylation, we combined reads from the positive and negative controls for methylation to obtain specific coverage (10x, 40x, 100x and 1000x) and percentages of methylation (0, 25, 50, 75, 100%) We then determined methylation using Nanopolish and the minimum coverage was defined when a significant difference was detectable in each group of percentage methylation (figure 1B) (Full list of p-values in supplementary table 1). Interestingly, even at the lowest level of coverage (10x) we observed a clear difference between the 0% and 75% and 100% methylation levels (p< 0.05), however, distinguishing between 25% and 50% was not possible at this level of coverage. At a coverage of 40x, a significant difference in 5mCpG levels between each percentage methylation group was observed, and as such, 40x was identified as the minimum sequencing depth required for detection of HBV methylation.

Starting with DNA extracted from HBV infected PHH or infected patient tissue, we utilized our cas9 enrichment protocol to linearise and sequence the HBV genome. The total yield ranged between 150-200K reads collecting in the range of 1-2 GB of DNA, our aligned read length was ∼2.5KB (N50 = ∼2520, median read length = ∼2310) (supplementary figure 1A). Raw reads were basecalled with Guppy (Version 4), a draft assembly was generated with Canu and polished with Medaka. The resulting consensus sequence was used to align the basecalled reads and calculate HBV genome coverage (figure 1C and 1D) and enrichment. We obtained a clear enrichment of HBV with up to 3000x coverage for certain HBV genotypes which, was comparable with total HBV detected in the same samples by qPCR and variability in the sequencing depth can explained by infection efficacy. Calculated with pycoQC (Leger and Leonardi 2019), ∼5% of all reads were on target (supplementary figure 1), which is a clear enrichment for this particular technique considering the very low levels of HBV detected with whole genome sequencing. Significantly, the gap present in the positive strand of rcDNA was identified in this analysis (supplementary figure 1). Levels of infection of the PHH used in this experiment were evaluated by qPCR, which established a high level of infection (figure 1C, 1D and supplementary figure 1). Given the high copy numbers of viral DNA, and thus replicative intermediates, it was expected to find this range in the length of the gap present in the positive strand of rcDNA, with a clear tapering of read length; this could be at least partially due to the different stages in the transcription of HBV DNA (supplementary figure 1). However, the lower coverage for the negative strand is likely due to the difference in efficacy of the sgRNA guides. Despite the differences in frequency, we were clearly able to sequence both strands of native HBV with cas9 enrichment technique at a coverage considered highly satisfactory for the identification of HBV methylation.

### Nanopore Flongles capable of sequencing native HBV in a mixture of viral and host DNA

In order to develop a more affordable and translatable method for the identification of HBV DNA in a mixture of viral and host DNA, we ambitioned to test our method on the smaller and cheaper Nanopore flow cells “Flongles”. Using these smaller flow cells allowed us to start with less DNA (0.5-1ug). This is an important consideration for clinical translation of this method since obtaining large quantities of DNA from tissue/ liquid biopsies is not be feasible. After 4h of sequencing we generated 367 reads totaling 1.3MB of sequencing data. 100 reads passed QC, 10% of which aligned to the HBV genome. However, we must note that the reads obtained from the flongle were of a much lower quality than those sequenced with the MinION flow cells (supplementary figure 2). While this was likely due to the lower molarity of DNA available for sequencing causing an increase in the speed at which DNA passes through the nanopores, causing a drop in translocation speed, it is, nonetheless, an important consideration for the potential applications of this technique. As expected, by using Flongles and starting with a lower quantity of starting material, we obtained lower coverage (∼10x) of the HBV genome than by using the larger and more expensive MinION flow cells (figure 1C). While our analysis to detect methylation differences indicated that 40x is the minimun required coverage to detect between 25% and 50% methylation, at a coverage of 10x we were able to significantly identify differences between 0% and 25% methylation, as well as 75% and 100% methylation. Thus a coverage of 10x for certain applications could be considered sufficient as is also the case for other techniques using short read sequencing (Olkhov-Mitsel and Bapat 2012) and long read sequencing (Gigante et al. 2019). Importantly, the same amount of starting material would have resulted in far greater coverage on the larger minION flow cell; however, while Nanopore is less expensive than other high throughput sequencing, the even lower price point of Flongle flow cells is an attractive alternative. Thus, our data suggests that our method can be applied to enrich HBV DNA in samples for sequencing with Nanopores ‘Flongles’ to obtain rapid detection of HBV in laboratory infection models.

**Figure 2.**
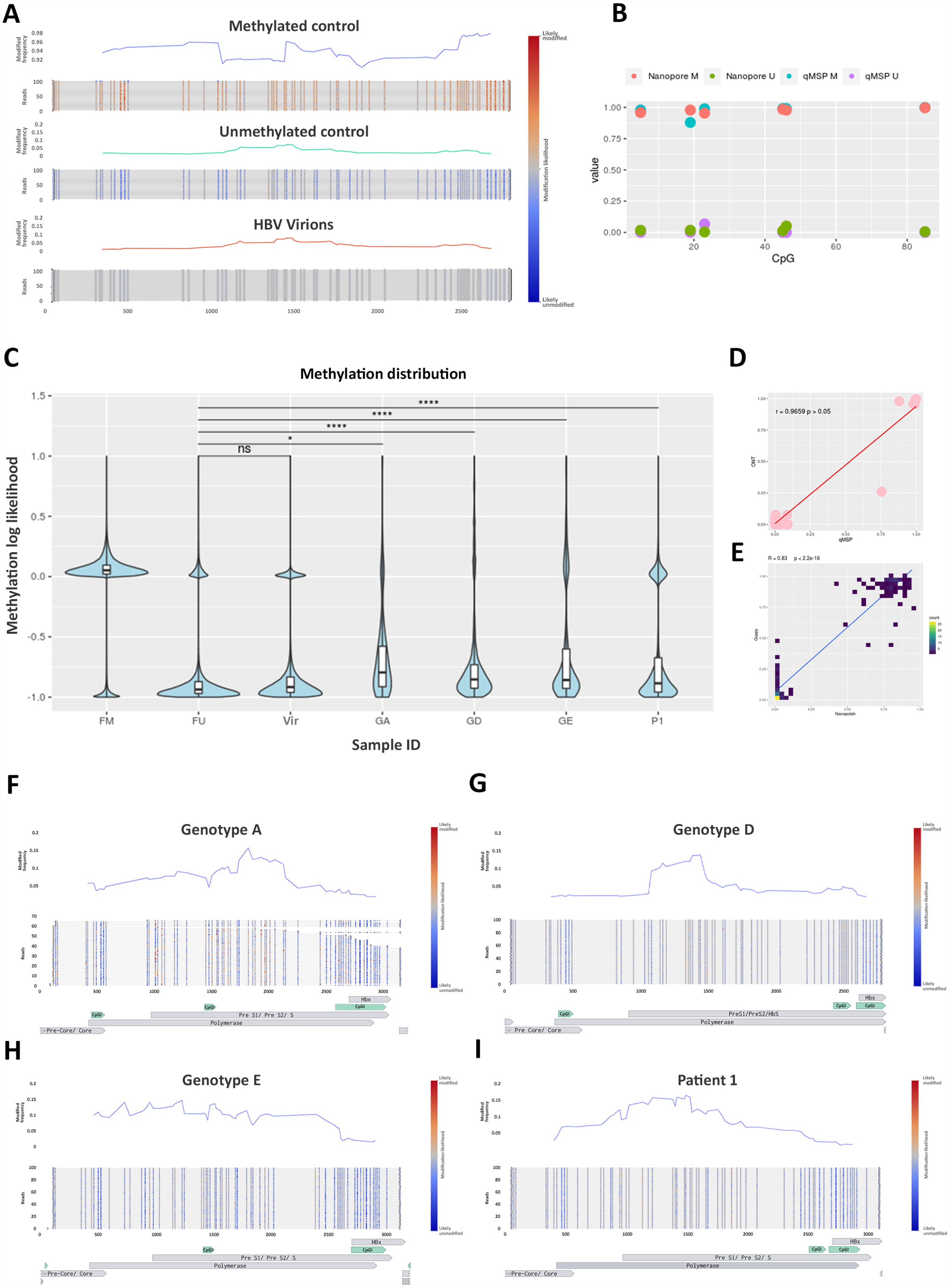
HBV methylation levels in HBV. **A:** Average methylation of HBV Controls and single molecule visualization of first 100 reads using methplotlib, **B:** 5mC levels of 6 CpG sites detected by 2 techniques; Nanopore sequencing and qMSP. **C:** Distribution of 5mC in HBV from PHH infected with different genotypes (GA, GD and GE) and isolated from patient tissue (P1). **D:** Correlation of 5mC levels obtained with qMSP with Nanopore, samples + controls. **E:** Correlation of 5mC levels detected with different methylation callers, Nanopolish and Guppy. **F-I:** Methylated frequency and single molecule visualization of HBV from infected PHH (**F**=Genotype A, **G**=Genotype D, **H**=Genotype E) and infected patient tissue (I=P1) (Blue = Unmethylted CpG site, Red = Methylated CpG site), (CpGI=CpG Islands detected with Meth primer).

### Nanopore sequencing detects HBV 5mCpG

We next sought to determine the validity of using Nanopores to detect modified bases in the HBV genome. Beta values for each CpG site were calculated by Nanopolish (Simpson et al. 2017) and plotted with methplotlib (De Coster, Stovner, and Strazisar 2020) (figure 2). Because nanopore sequencing directly evaluates methylation patterns on native DNA strands, we are able to observe long-range methylation information on each DNA molecule. As anticipated, very low levels of methylation were observed in the negative control with some residual methylation and background noise detected (figure 2A). Interestingly, we identified some single reads that were methylated in the negative control, likely a result of the presence of residual un-amplified DNA; thus, we determined that low levels of methylation observed were partially attributed to this contaminating starting material. However, we also identified random methylated CpG sites throughout reads in the fully unmethylated control. These could also be due to methylation calling errors and can therefore be considered as background noise for the technique. In the positive control we detected high levels of DNA methylation (figure 2A). However, there was variability in the average levels of 5mCpG. After visualising the methylation of the single molecules, we identified a number of reads that were not fully methylated which was likely contributing to the lower average levels of 5mCpG at certain loci. These unmethylated reads are likely attributed to the efficiency of the Methyltransferase in the preparation step of the fully methylated control. However, in a similar way to the unmethylated control, there was also some random CpG sites throughout the data that were not methylated in the positive control, which is more likely attributed to methylation calling errors. In order to address the issue of introduced bias through methylation callers, in addition to calling 5mCpG with Nanopolish, we validated our findings with an additional methylation caller, Guppy. Spearmans correlation of the methylation levels obtained for the fully methylated and unmethylated HBV controls with Nanopolish and Guppy + Medaka, indicated a highly significant correlation (R = 0.83, p< 2.2e^-16^) (figure 2E).

In order to validate our findings obtained with Nanopore, we developed a bisulfite quantitative methyl-specific qPCR assay (BS-qMSP). Briefly, DNA from controls and genotype D samples were bisulfite converted, followed by methyl-specific qPCR, with methyl specific primers designed for 6 CpG sites in the HBV genome. Beta Values were calculated by comparing the % of methylated DNA to the total (unmethylated + methylated DNA). (Figure 2B). As expected, beta values were comparable for all CpG sites in the methylated, unmethylated controls and samples, and exhibited a high correlation coefficient (0.96, P< 0.05) (figure 2D).

The clear extremes observed in the different controls (figure 2A) (> ∼90% methylation in FM, < ∼10% methylation in the FU) validated in multiple methylation callers (figure 2E), and with an additional technique (figure 2B, 2D), indicate the efficacy of Nanopore technology for the detection of 5mCpG on HBV DNA. We were therefore highly confident in this tool for the detection of 5mCpG levels on the HBV genome.

### Absence of DNA methylation observed in HBV virions

After identifying the background noise levels of 5mCpG identified in synthetically fully methylated and unmethylated HBV DNA, we next sought to determined the 5mCpG levels in an expected biologically negative control, HBV Virions. HBV DNA is reverse transcribed after being packaged in the viral capsid in the cytosol of the infected cells, thus out of touch with DNA methyltransferase enzymes and as a result, likely not methylated (Wang and Seeger 1993). Overall, there were very low levels of 5mCpG identified in the HBV virions, comparable to the amplified HBV control (figure 2A). These levels were not higher than those detected in the unmethylated HBV control (figure 2C), and as such, were likely a result of methylation calling errors or potentially some contaminating DNA from dead cells also collected during the HBV viral particle purification process. Regardless, the levels were still very low (< 15%). Taken together with the methylation levels in the amplified, unmethylated HBV control, these levels of 5mCpG indicate the background noise levels of the detection method. Moreover, these data continued to increase our confidence in Nanopore sequencing to accurately detect 5mCpG in the HBV genome.

### Basal levels of HBV 5mCpG in infected PHH with different genotypes

In order to determine the basal levels of 5mCpG in HBV infection models in vitro, we used primary human hepatocytes (PHH) infection model, which is the gold standard HBV in vitro infection model. PHH were infected with a viral inoculum of HBV genotypes A, D and E and collected nuclear DNA from cells 6 days post infection. Infection efficiency was determined by HBsAg and HBeAg expression by ELISA as well as total HBV DNA in the supernatant by qPCR every 3 days. We observed similar infection rates for HBV Genotype D and E, however, for genotype A all parameters were negligible, indicating lower infection rates (supplementary figure 1). Despite the negligible levels detected for the infection efficacy of HBV genotype A, our cas9 enrichment technique was able to enrich and sequence HBV genotype A with ∼80x coverage (figure 1C). In addition, our cas9 technique was highly effective at enriching HBV of all genotypes tested at sufficient coverage to evaluate methylation levels.

Detected 5mCpG levels were comparable across the different HBV genotypes (figure 2F-H). We observed low levels of 5mCpG across the three genotypes tested with HBV genotype E having the highest basal levels across the genome (figure 2C). While we did not identify any differentially methylated loci when comparing the different genotypes, the technique developed was clearly able to enrich and sequence all genotypes tested. Moreover, the distribution of methylation for each of the different genotypes was significantly different from both of the negative controls (figure 2C) and from each other. Interestingly we also identified an enrichment of 5mC in the preS1/ preS2 region for each of the genotypes and could thus have implications for HBS expression. These differences in 5mC distribution taken together with the different pattern of methylation frequency (figure 2F-H) indicate that HBV genotypes exhibit genotype specific 5mC landscapes. Thus, these data further add to the efficacy of the cas9 sequencing technique for the enrichment and delineation of HBV methylation in a laboratory setting.

### Enrichment and sequencing of HBV in infected patients identifies epigenetic heterogeneity

In order to test the efficacy of our technique and its potential for translational research in clinics, we proceed to test cas9 enrichment and sequencing in HBV DNA from patient tissue. We took advantage of samples collected as part of the PROLIFICA study (Lemoine et al. 2016). We used 900ng of DNA obtained from a HBV positive patient’s liver biopsy. All viral parameters were assessed including pgRNA and quantification of cccDNA (supplementary Table 2). By using our cas9 sequencing technique we were able to enrich in enough HBV specific reads to perform a *de novo* assembly. The HBV genotype was determined by jumping profile Hidden Markov Model (jpHMM) and identified as genotype A, and after alignment, identified over 2000x coverage of this HBV genome (figure 1C and 1D). The methylation of CpG sites was determined and the average 5mCpG levels were calculated (figure 2I). Interestingly, certain CpG sites were around 50% methylated across the HBV genome. Visualization of the single HBV DNA molecules revealed heterogeneity in the methylation levels, indicating the potential existence of differentially methylated HBV populations within the patient (figure 3). In order to explore this data further, we evaluated the distribution of HBV methylation levels in the patient sample. We observed a significant difference in the distribution of 5mCpG compared to both the fully methylated and fully unmethylated controls (figure 2C). Visualization of the 5mCpG distribution of individual reads (figure 3A) revealed heterogeneity of HBV molecules. Unsupervised hierarchical clustering of whole HBV molecules, identified 4 distinct clusters (figure 3B-C), suggesting that within the 1 patient, up to 4 epigenetically distinct HBV phenotypes exist. Whilst the differences between clusters was not identified as a particular region, there were certainly several CpGs that displayed large differences between groups. In particular CpG in the preS1/ preS2 S region that displayed the large differences between clusters (figure 3D and 3E). While, these data need to be confirmed with additional patients to conclude more broadly, it is not entirely surprising; HBV replication intermediates and virions were not methylated, and the methylation levels observed are likely due to the presence of cccDNA. An understanding of the epigenetic landscape of different HBV DNA populations is important when considering HBV regulation and occult infection. Taken together, these data highlight the potential of this technique in the enrichment and sequencing of HBV as well as analysis of HBV DNA methylation in clinical settings.

**Figure 3.**
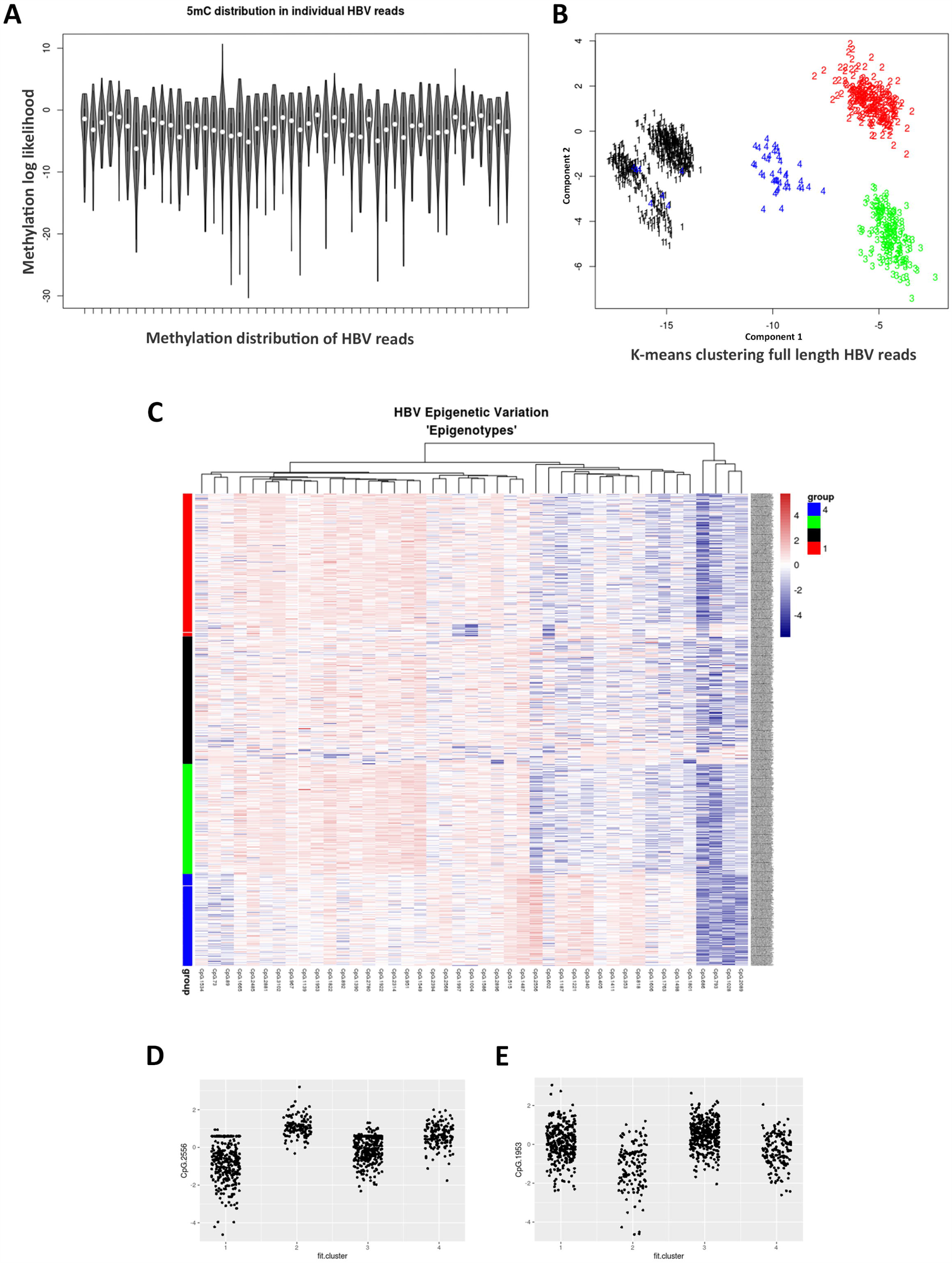
Epigenetic heterogeneity in HBV from infected patients. **A:** Variability of 5mC levels in a random selection of single HBV molecules. **B:** K-means clustering of HBV molecules. **C:** heatmap clustering of single HBV molecules.

## Discussion

Mapping the methylation of HBV DNA in infected cells can alter the expression patterns of viral genes related to infection and cellular transformation (Fernandez et al. 2009; Guo et al. 2009) and may also aid in understanding why certain infections are cleared, or persist with or without progression to cancer (Mirabello et al. 2012). Furthermore, the clear detection of viral methylation patterns could potentially serve as biomarkers for diseases that are currently lacking, including occult HBV infection (Nakamura et al. 2020). However, development of more sensitive high throughput techniques translatable to the clinic, is essential. The present study developed a technique to enrich and sequence HBV without the need for bisulfite conversion or PCR using cas9 guided RNPs coupled with Nanopore sequencing. We were able to enrich and sequence HBV from both infected PHH and patient liver tissue achieving coverage of ∼2000x providing the first *de novo* assembly of native HBV DNA, as well as the first landscape of 5mCpG from native HBV sequences. By using nanopore we also took advantage of sequencing entire HBV genomes and identified several HBV epigenotypes in patient tissue.

The cas9 enrichment technique was first designed to enrich in target regions in the human genome prior to sequencing (Gilpatrick et al. 2020). We have adapted this method to enrich in viral DNA in a mixture of host DNA. The coverage obtained for the previous study was ∼400x by using a triple cutting approach, whereby six or more sgRNAs were designed for each region of interest. By using just 2 sgRNAs targeting the HBV genome, we obtained much higher coverage than in the previous study and dramatically reduced the cost/ sample. We have attributed this to the smaller size of the HBV genome compared to the average sized regions the authors were targeting (3.2kb v 20kb) as well as the higher concentration of HBV genomes/ cell. Thus, it is not entirely surprising that the coverage was improved by such a large factor (>25x).

The developed technique enriches for episomal HBV DNA and can also somewhat distinguish between the different HBV forms. We first blocked linear DNA ends by phosphorylation and then linearized HBV with cas9 guided RNPs, after sequencing we selected reads that were less than 4000bp to perform a de novo assembly. In doing so, we were sure to remove any integrated HBV reads that were sequenced, increasing specificity for episomal HBV. Moreover, we identified the “gap” in the positive strand of HBV. Therefore, during post-sequencing analysis, by selecting only HBV reads that cover the whole genome we identified those reads that correspond to cccDNA’s positive strand. However, since there is no gap in the negative strand of HBV rcDNA we can not distinguish it from cccDNA in this way. Other techniques to study rcDNA and cccDNA are limited. Southern Blot remains the gold standard for quantification of cccDNA, however it is time consuming and not practical for a large number of samples (Xia et al. 2017). Other techniques rely on PCR but show limited specificity when excess replication intermediates are present. Furthermore, HBV is renowned for its genomic sequence variability, with different genotypes varying as much as 7.5% (Rajoriya et al. 2017) which can prove difficult for the design of primers. Our cas9 enrichment protocol was capable of sequencing all HBV genotypes tested. This is likely due to the conservation of the region used for the guides designed to specifically target HBV DNA which was not bound by polymerase kinetics. Our cas9 technique was also capable of enriching in HBV from patient liver tissue achieving over 2000x coverage of the HBV genome and thus demonstrating the potential of this technique in translation to clinical applications.

Current strategies for DNA methylation determination include bisulfite conversion followed by Sanger sequencing or methyl specific PCR, enzyme digestion assays and single-molecule, real-time (SMRT) sequencing (Olkhov-Mitsel and Bapat 2012). It is difficult to compare the efficiency of different techniques for detecting DNA methylation, since they are all valuable for different applications; despite this, thorough reviews and comparisons have been conducted previously with limitations for each platform well described (Olkhov-Mitsel and Bapat 2012). Our technique was capable of detecting 5mCpG in native HBV DNA from different genotypes in vitro, as well as in patient liver tissue. An advantage of our cas9 enrichment technique compared to traditional biulfite-based sequencing methods is the clear benefit of sequencing native DNA that does not require Bisulfite conversion. Bisulfite conversion results in DNA degradation and, importantly, is not capable of distinguishing 5mC from other modified bases such as 5hmC, whereas Nanopore based techniques are able to do so. In addition, with this nanopore technique we are able to simultaneously elucidate the DNA sequence and the single molecule methylation, while other techniques like Bisulfite conversion followed by methyl specific PCR require additional experimental analysis to ascertain the sequence information. Furthermore, by sequencing full HBV reads we are able to identify the heterogeneity of HBV sequences making it possible to identify certain reads that are displaying differential methylation patters within the sample population, another application that is not possible with existing techniques. The exception of course is SMRT sequencing. However, this technique has prohibitively expensive set up and ongoing costs compared to Nanopore, making it unfeasible for rapid deployment and set up of viral monitoring potentially in remote locations. In comparison, Nanopore is light, easy to handle and by using Fongles we obtained whole genome length HBV sequences in < 4h.

While the nanopore platform has clear benefits there are also several limitations. Importantly, a large quantity of starting material is required for this technique (1-5µg of DNA). While this is true on paper, we were able to achieve 2000x coverage of HBV with 900ng of starting material (liver tissue from a HBV infected patient). On average our Patient liver tissue biopsies are less than 25mg and depending on the DNA extraction technique we can obtain up to 1-2µg of DNA. However, for other patient samples such as blood and plasma it is simply not possible to obtain large quantities of DNA. As such, PCR and bisulfite-based methods must be preferred. Nevertheless, with the development of the smaller nanopore Flongles and the continually improving bioinformatics and machine learning applications (Payne et al. 2020) it will soon be feasible to combine our technique with bioinformatic approaches to enrich in viral DNA from much lower quantities of starting material.

While we have focused on the application of this technique for detecting modified viral DNA bases, this tool can also be useful for other approaches; including the identification of mutations and tracking viral evolution, investigate transmission chains, the likes of which were observed in the Ebola outbreak in West Africa (Quick et al. 2016) and more recently in the COVID-19 pandemic of 2020 (Viehweger et al. 2019). By sequencing native reads and performing a *de novo* assembly, introduced bias from PCR amplification is eliminated. However, nanopore sequencing is still considered error prone with the raw read accuracy varying depending on several factors such as flow cell type, starting material quality, basecaller, basecalling speed, polishing etc. In optimal conditions, nanopore sequencing has a reported raw read accuracy of ∼97% using flow cells with the R9.4 pore chemistry, while, the consensus accuracy and single molecule consensus accuracy is usually around 99.98% (Q37) (Lu, Giordano, and Ning 2016). In the present study, our *de novo* assembly of native HBV reads generated a consensus with per-base 99.9% similarity to existing HBV genotype D references generated with illumina sequencing (supplementary figure 1C), which is comparable to the benchmark values for the assembler used (Canu) (Koren et al. 2017; Nurk et al. 2020). Accurate references are imperative to improve the accuracy of downstream analysis, such as methylation calling.

Furthermore, there is huge potential for nanopore technology and the identification of additional modified bases. Since only low levels were observed and only 1 patient biopsy was screened in the present study, we were not able to correlate methylation with functional outcomes such as expression of viral parameters. That being said, the study of epigenetics is still in its infancy. Recently we found that levels of 5hmC in gene bodies in differentiating hepatocytes is highly correlated with the expression of those genes (Rodríguez-Aguilera et al. 2019). Thus, there is still a lot more to discover regarding the function of modified bases. However, the first point of call is accurate detection, and more work is needed to develop machine learning tools to detect, and benchmark additional modified bases in native Nanopore data.

## Conclusions

We developed a sensitive and high throughput method for the enrichment and nanopore sequencing of native HBV DNA from infected PHH and patient liver tissue. This method is a novel approach that achieved a clear enrichment of viral DNA in a mixture of virus and host DNA without the need for PCR amplification. By sequencing native HBV DNA with nanopore we were also able determine the DNA methylation landscape of HBV without the use of bisulfite conversion. Moreover, using the developed technique, we have provided the first *de novo* assembly of native HBV DNA, as well as the first landscape of 5mCpG from Naive HBV DNA. More work is needed to test compatible machine learning enrichment techniques to further improve the minimum required concentration of starting material.

## Methods

### Cultivation of PHH

Primary Human Hepatocytes (PHH) were extracted and maintained as previously described (Ancey et al. 2015).

### HBV cultivation and infections

HBV inocula was generated as previously described (Ancey et al. 2015; Lucifora et al. 2011). PHH were naturally infected with HBV genotype A, D and E for 24h (MOI 100). A stable infection was achieved after 3 days, cells were maintained for up to 14 days. Infection efficacy was determined by quantification of Hepatitis B surface antigen (HBsAg) and Hepatitis B e antigen (HBeAg) concentration in supernatant by ELISA and calculation of HBV copies/µL by qPCR as previously described (Ancey et al. 2015).

### Patient liver tissue

Patient samples were collected as a part of the PROLIFICA study (Cohen et al. 2020; Lemoine et al. 2016). DNA was extracted from snap frozen human liver needle biopsies. Liver samples were first homogenized on ice using a TissueRuptor (Qiagen, Hilden, Germany) in homogenization buffer (Tris HCl pH 8, 50 mM; EDTA 1 mM; NaCl 150 mM) and then processed for DNA extraction.

### DNA extraction

Cells, or homogenized tissues were digested with proteinase K prior to DNA isolation using MasterPure™ DNA Purification Kit (Epicentre, by Illumina, Madison, United States) according to manufacturer’s instructions.

### Quantification of total HBV-DNA, cccDNA and pregenomic(pg) RNA in liver sample

Quantification was performed using the QX200^™^ Droplet Digital PCR System (BioRad, Hercules, California, USA) with primers and fluorescence dual hybridization probes specific fortotal HBV-DNA or cccDNA as described in (Lebossé et al. 2017; Testoni et al. 2019). Before cccDNA amplification, DNA was treated with 10U of Plasmid-safe Dnase (Epicentre^®^ Illumina) for 45 minutes at 37°C following thelatest update of the international working group on cccDNAstandardization (Allweisset al., 2017 International HBV Meet-ing, O-45). Serial dilutions of a plasmid containing an HBVmonomer (pHBV-EcoR1) served as quantification standard. To normalize the number of viral copies per cell content, the number of cellular genomes was determined using the b-globin genekit (Roche Diagnostics, Manheim, Germany). Patient sample was independently analyzed in duplicate. The range of quantification was comprised between 10^1^ and 10^7^ copies of HBV genome/well for both cccDNA and total HBV-DNA assays. For pgRNA detection, specific primers and Taqman^®^ hybridization probe were used, as described in (Lebossé et al. 2017; Testoni et al. 2019). The patient sample was independently analyzed in duplicate and pgRNA relative amount was normalized over the expression of housekeeping the gene GUSB (Hs99999908_m1, Thermo Fisher Sci-entific, Waltham, MA, USA).

### Laboratory assays

HBsAg was detected by chemiluminescent microparticle immunoassay (Architect, Abbott) in 2012–2013 (Njai et al. 2015). HBV DNA levels were measured at the end of the Prolifica study in stored serum samples using in-house quantitative real-time PCR (detection limit: 50 IU/mL), calibrated against an international standard (Ghosh, Sengupta, and Scaria 2014).

### Fully unmethylated and fully methylated controls

HBV DNA was amplified by nested PCR as previously described to prepare a negative (fully unmethylated) control. After amplification, a positive control for methylation (fully methylated) was prepared by methylating CpG dinucleotides; by incubating 1µg of DNA with S-Adenosyl methionine (SAM) (32µM) with CpG Methyltransferase (M.SssI) (4-25 units) (New England BioLabs) at 37°C for 1h before heating to 65°C for 20mins.

### Nanopore library prep and sequencing

DNA (0.5-3µg) from each sample or control was enriched in HBV and linearized using cas9 guides RNPs. TracrRNA and crRNA (5’AGCTTGGAGGCTTGAACAGT3’ and 5’TAAAGAATTTGGAGCTACTGTG3’) were purchased from Integrated DNA Technologies (IDT). Samples were barcoded and multiplexed using the Nanopore Ligation Sequencing kit (SQK-LSK109) and Native barcode expansion kit according to manufacturer’s instructions (Oxford Nanopore Technology, Oxford UK). Sequencing was conducted with a Minion sequencer on ONT 1D flow cells (FLO-min106) with protein pore R9.4 1D chemistry for 24h or on a Flongle (FLO-106) for 4h Oxford Nanopore Technology, Oxford UK). Reads were basecalled with Guppy (version 4).

### De novo assembly

Basecalled fasQ files were used to assemble the HBV genome with canu (Koren et al. 2017), that was polished using Medaka; a tool to create a consensus sequence from nanopore sequencing data using neural networks applied from a pileup of individual sequencing reads against a draft assembly. Basecalled FastQs were then aligned to the generated consensus sequence using Minimap2 (Li 2018). Assembly was assessed by using BLAST (Zhang et al. 2000) which returned a 99.9% similarity score to existing HBV genotype D references and jumping profile Hidden Markov Model (jpHMM) which correctly identified the genotypes (supplementary figure 1C).

### Methylation calling

We first determined the methylation status of each CpG site on every read by using the widely used tool, nanopolish (Simpson et al. 2017) used recently by (Gigante et al. 2019). For validation, we also called DNA methylation using novel tool, Guppy (version 4). CpG Islands were predicted using MethPrimer.

### Bisulfite (BS) and quantitative methyl specific PCR (qMSP)

BS and qMSP protocols made available as detailed methods at protocols.io (Hernandez and Goldsmith n.d.). Primer sequences are available in supplementary table 3. Primers were designed with MethPrimer and purchased from Thermo Fisher Sci-entific (Waltham, MA, USA).

### Statistics

Data processing and statistic analyses were performed in R bioconductor. Differential methylation was determined by Dispersion shrinkage for sequencing data (DSS) as previously described (Gigante et al. 2019; Park and Wu 2016). Kruskal–Wallis one-way analysis of variance was used to determine differences in average methylation levels. Significance value = p< 0.01. Hierarchical clustering was performed by k-means cluster analysis using d-Ward Hierarchical clustering with the elucidean method, number of groups determined by the elbow method.

### Converting reads for visualization in IGV

In order to take advantage of the single molecule sequencing with Nanopore, we converted the reads for visualization using Methplotlib (De Coster et al. 2020).

## Data availability

All raw and processed sequencing data is has been made publicly available with GEO (Ascension numbers: GSE162518, GSM4953612, GSM4953613, GSM4953614, GSM4953615, GSM4953616, GSM4953617).

Assembled HBV sequences made available in GeneBank Submission ID # 2386637.

### Abbreviations list

5mC: 5-methyl cytosine
5mCpG: 5-methyl cytosine in CpG context
RNP: Riboneucleotide protein-primed
HCC: Hepatocellular carcinoma
pgRNA: Pregenomic RNA
rcDNA: Relaxed circular DNA
cccDNA: Covalently closed circular DNA
HBV: Hepatitis B Virus
5hmC: 5-hydroxy methyl cytosines
PCR: Polymerase chain reaction
PHH: Primary Human Hepatocytes
GA: Genotype A
GD: Genotype D
GE: Genotype E
BS-qMSP: Bisulfite quantitative methyl-specific qPCR
FU: Fully Unmethylated Controls
FM: Fully Methylated Controls

## Conflict of interest

Chloe Goldsmith and Hector Hernandez have received travel and accommodation support to attend conferences for Oxford Nanopore Technology.

## Acknowledgements

The Authors would like to thank the patients that participated in this study and also acknowledge the whole Inserm U1052 team for any help along the way.

## Funding

This work was supported by the Agence Nationale de Recherches sur le SIDA et les Hépatites Virales (ANRS, Reference No. ECTZ47287 and ECTZ50137); La Ligue

Nationale Contre Le Cancer Comité d’Auvergne-Rhône-Alpes AAP 2018.

### Ethics approval and consent to participate

Pre-test counseling was delivered and written consent obtained. Ethics approval for the study was granted by the Government of The Gambia and MRC Gambia Joint Ethics Committee.

## Authors’ contributions

CG generated concepts, completed experiments, performed analysis and wrote the manuscript: DC and AD completed experiments: GM assisted in concept generation and manuscript preparation: BT assisted in manuscript preparation: KP assisted in manuscript preparation: AC obtained funding and assisted in manuscript preparation: HH performed analysis and obtained funding: IC assisted in manuscript preparation and obtained funding. All authors have discussed the results and read and approved the manuscript.

## Supplementary Figure Legends

**Supplementary figure 1**. A and B: Yield for aligned reads calculated with pycoQC (representative sample). C: Coverage of HBV identifying the Gap in HBV reads. D: HBV genotype was determined by jumping profile Hidden Markov Model (jpHMM). E-F: Infection efficacy for HBV genotypes. PHH were infected with 100MOI of HBV geotype A, E or D. Serum levels of HBsAg, HBeAg were quantified by ELISA and HBV DNA was quantified by qPCR after 0, 3 and 6 days post infection.

**Supplementary figure 2**. PHRED scores for nanopore runs. A: Genotype A sequenced on MinION, B: Genotype D sequenced on MinION, C: Genotype E sequenced on MinION, D: Genptype D sequenced on Flongle.

**Supplementary Table Legends**

**Supplementary table 1**. Sequencing depth for effective calculation of HBV 5mCpG levels: p-value table corresponding to figure 1B.

**Supplementary table 2**. Primer sequences for BS-qMSP of HBV.

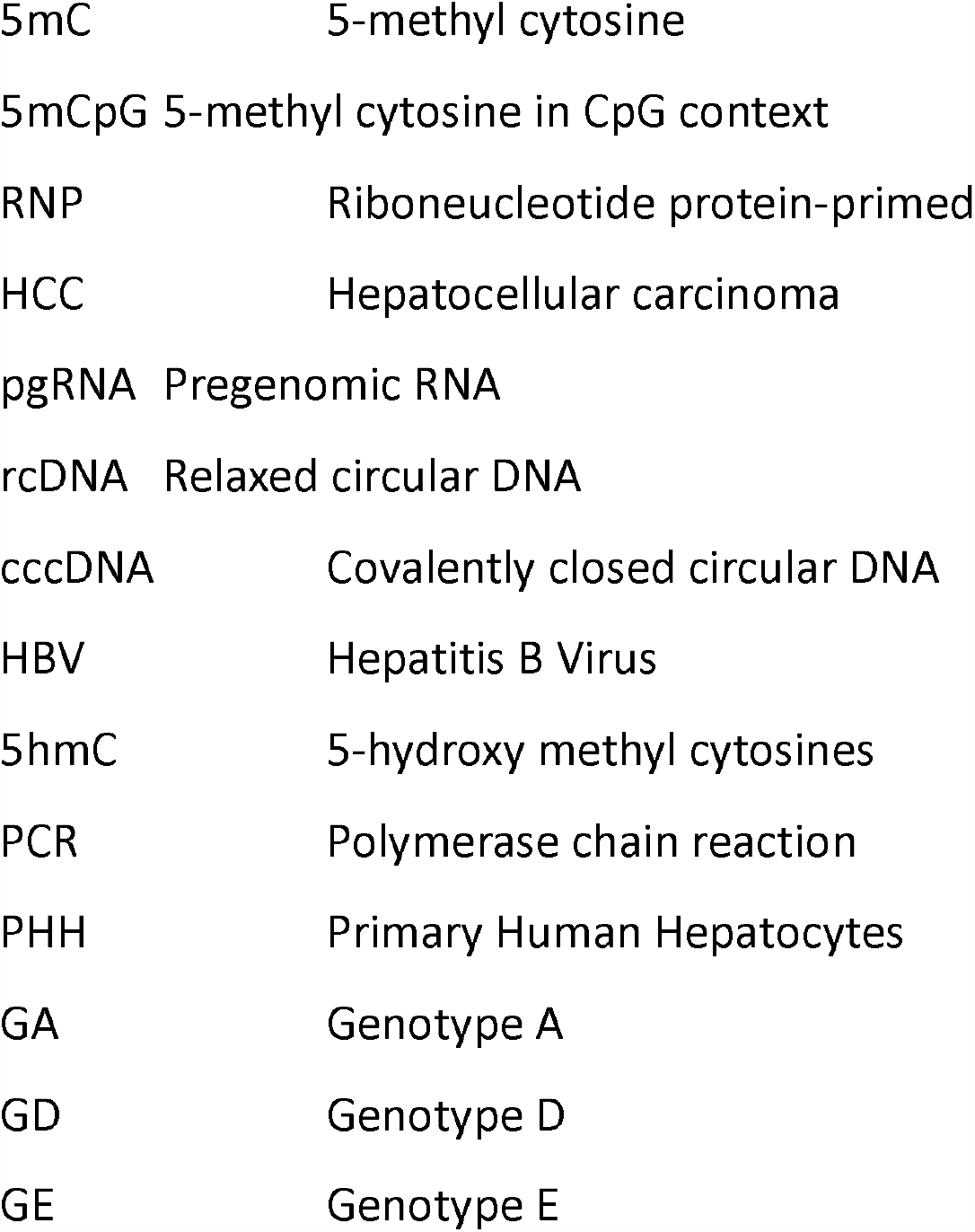
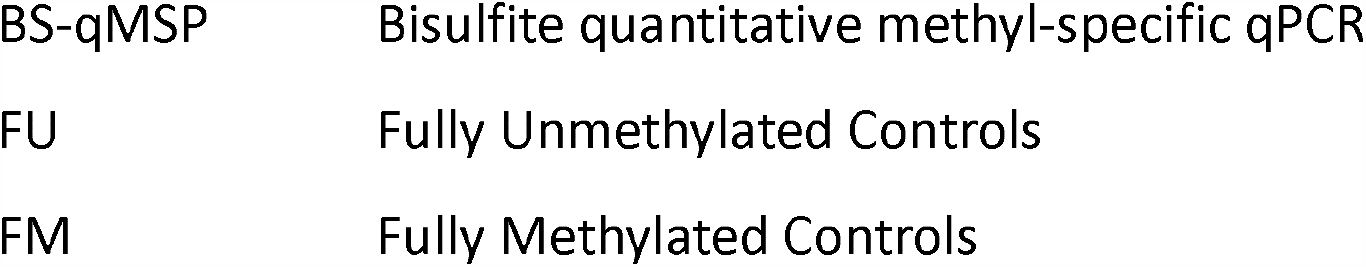

## Notes

### Summary of Updates

Updated Abstract

