## Supplementary figures and images for "Epigenetic heterogeneity after *de novo* assembly of native full-length Hepatitis B Virus genomes"

### Supplementary Figure 1

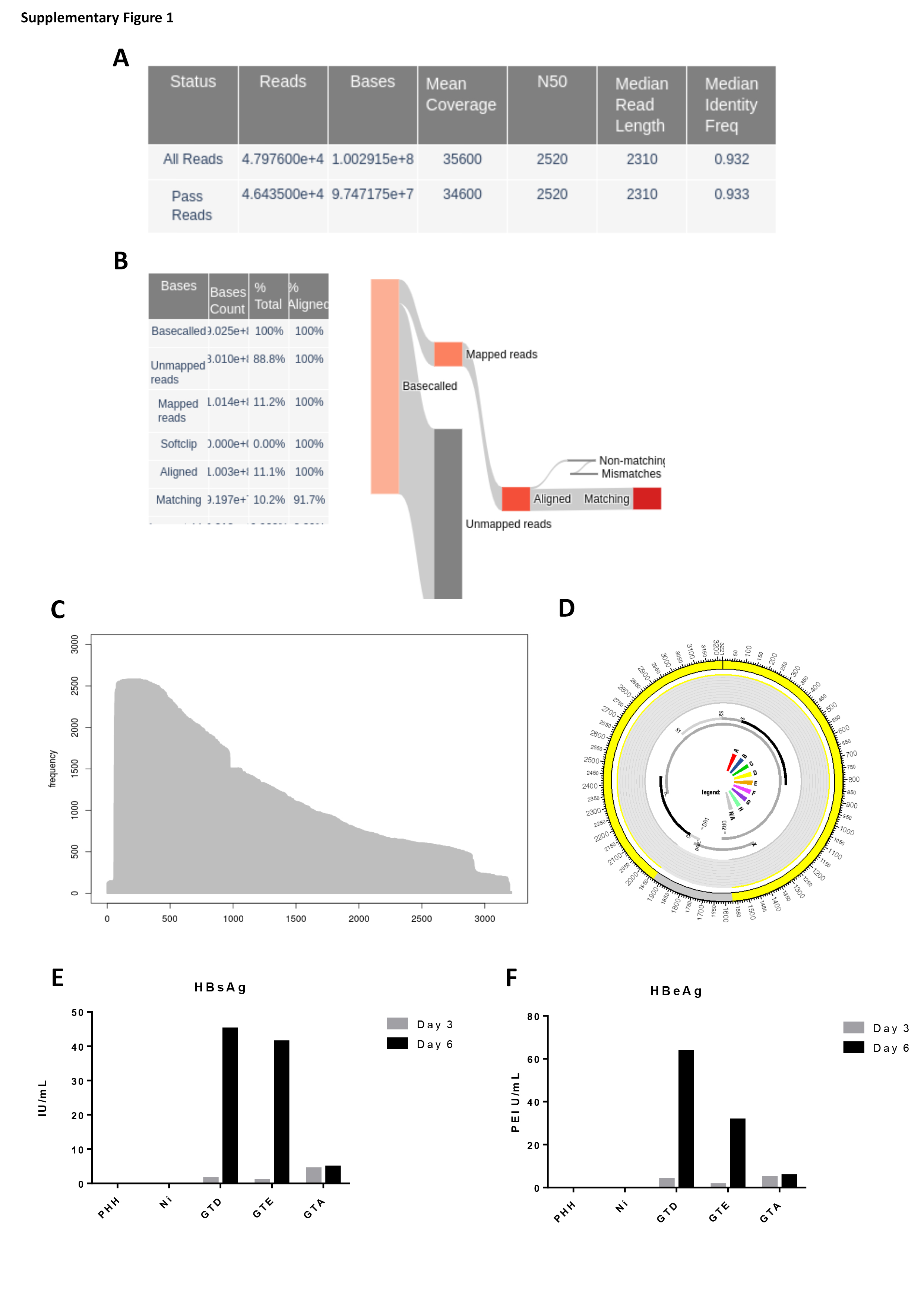

### Supplementary figure 2

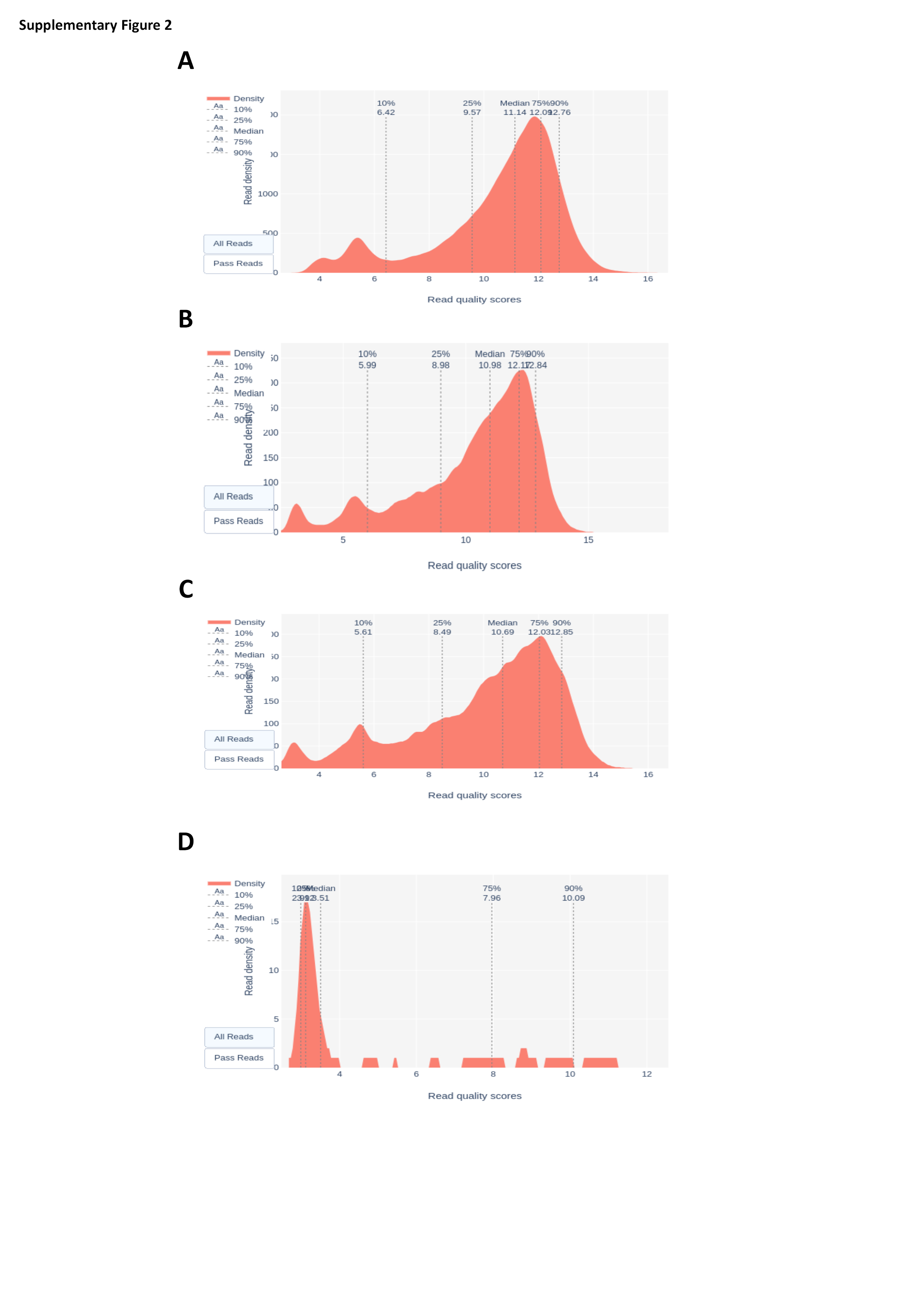
