## Supplementary tables 1 and 2 for "Epigenetic heterogeneity after *de novo* assembly of native full-length Hepatitis B Virus genomes"

**Supplementary table 1**

| coverage | .y. | Group 1 | Group 2 | statistic | p | padj | p.adj.significant |
|---|---|---|---|---|---|---|---|
| **10** | Methylation | 1_0M | 2_25M | 527.5 | 2.05E-12 | 8.2E-11 | **** |
| **10** | Methylation | 1_0M | 3_50M | 392 | 1.44E-14 | 5.76E-13 | **** |
| **10** | Methylation | 1_0M | 4_75M | 103 | 3.21E-20 | 1.284E-18 | **** |
| **10** | Methylation | 1_0M | 5_100M | 69 | 7.03E-23 | 2.812E-21 | **** |
| **10** | Methylation | 2_25M | 3_50M | 1391.5 | 0.043 | 1 | ns |
| **10** | Methylation | 2_25M | 4_75M | 485.5 | 4.59E-12 | 1.836E-10 | **** |
| **10** | Methylation | 2_25M | 5_100M | 168 | 2.41E-19 | 9.64E-18 | **** |
| **10** | Methylation | 3_50M | 4_75M | 647.5 | 2.13E-09 | 8.52E-08 | **** |
| **10** | Methylation | 3_50M | 5_100M | 272 | 2.94E-17 | 1.176E-15 | **** |
| **10** | Methylation | 4_75M | 5_100M | 342 | 5.43E-16 | 2.172E-14 | **** |
| **40** | Methylation | 1_0M | 2_25M | 193 | 2.06E-17 | 8.24E-16 | **** |
| **40** | Methylation | 1_0M | 3_50M | 127.5 | 1.42E-18 | 5.68E-17 | **** |
| **40** | Methylation | 1_0M | 4_75M | 58 | 3.38E-20 | 1.352E-18 | **** |
| **40** | Methylation | 1_0M | 5_100M | 1.5 | 7.56E-22 | 3.024E-20 | **** |
| **40** | Methylation | 2_25M | 3_50M | 1148 | 0.00095 | 0.038 | * |
| **40** | Methylation | 2_25M | 4_75M | 223 | 1.27E-16 | 5.08E-15 | **** |
| **40** | Methylation | 2_25M | 5_100M | 55.5 | 2.2E-20 | 8.8E-19 | **** |
| **40** | Methylation | 3_50M | 4_75M | 335 | 2.4E-14 | 9.6E-13 | **** |
| **40** | Methylation | 3_50M | 5_100M | 104 | 3.29E-19 | 1.316E-17 | **** |
| **40** | Methylation | 4_75M | 5_100M | 180 | 7.93E-18 | 3.172E-16 | **** |
| **100** | Methylation | 1_0M | 2_25M | 306.5 | 6.15E-15 | 2.46E-13 | **** |
| **100** | Methylation | 1_0M | 3_50M | 80 | 3.13E-19 | 1.252E-17 | **** |
| **100** | Methylation | 1_0M | 4_75M | 60 | 7.82E-20 | 3.128E-18 | **** |
| **100** | Methylation | 1_0M | 5_100M | 8 | 4.73E-21 | 1.892E-19 | **** |
| **100** | Methylation | 2_25M | 3_50M | 352.5 | 5.01E-14 | 2.004E-12 | **** |
| **100** | Methylation | 2_25M | 4_75M | 82 | 1.96E-19 | 7.84E-18 | **** |
| **100** | Methylation | 2_25M | 5_100M | 8 | 4.29E-21 | 1.716E-19 | **** |
| **100** | Methylation | 3_50M | 4_75M | 340 | 2.99E-14 | 1.196E-12 | **** |
| **100** | Methylation | 3_50M | 5_100M | 33.5 | 2.27E-20 | 9.08E-19 | **** |
| **100** | Methylation | 4_75M | 5_100M | 139.5 | 2.39E-18 | 9.56E-17 | **** |
| **1000** | Methylation | 1_0M | 2_25M | 41.5 | 4.12E-20 | 1.648E-18 | **** |
| **1000** | Methylation | 1_0M | 3_50M | 1 | 7.94E-21 | 3.176E-19 | **** |
| **1000** | Methylation | 1_0M | 4_75M | 0 | 5.18E-21 | 2.072E-19 | **** |
| **1000** | Methylation | 1_0M | 5_100M | 0 | 5.15E-21 | 2.06E-19 | **** |
| **1000** | Methylation | 2_25M | 3_50M | 232 | 3.03E-16 | 1.212E-14 | **** |
| **1000** | Methylation | 2_25M | 4_75M | 68 | 1.01E-19 | 4.04E-18 | **** |
| **1000** | Methylation | 2_25M | 5_100M | 4 | 4.31E-21 | 1.724E-19 | **** |
| **1000** | Methylation | 3_50M | 4_75M | 244 | 5.14E-16 | 2.056E-14 | **** |
| **1000** | Methylation | 3_50M | 5_100M | 18.5 | 1.31E-20 | 5.24E-19 | **** |
| **1000** | Methylation | 4_75M | 5_100M | 93.5 | 3.4E-19 | 1.36E-17 | **** |

**Supplementary table 2**

| NAME | SEQUENCE (5' > 3')<br>5-139 bases according to the synthesis scale |
|---|---|
| CpG1MF1 | ATTTTTGGAAGAGAAATCGTTA |
| CpG1MR1 | CGTCGTCTAACAACAATAATTTCC |
| CpG1UF1 | ATTTTTGGAAGAGAAATTGT |
| CpG1UR1 | AAAAAACCTACCTCATCATCT |
| CpG2MF1 | TATATATTTTATGGAAGGCGGG |
| CpG2MR1 | AACTAAATCCAACTAATAATCGAAA |
| CpG2UF1 | ATATATTTTATGGAAGGTGGG |
| CpG2UR1 | ACTCTAAAAACTAAATCCAACTAATAATCA |
| CpG3MF1 | TAATAAGGTAGGAGTTGGAGTATTCG |
| CpG3MR1 | ATTTCTCAAAAATAAAAACAACGAAA |
| CpG3UF1 | AATAAGGTAGGAGTTGGAGTATTT |
| CpG3UR1 | ATTTCTCAAAAATAAAAACAACAAAA |
| CpG4MF1 | TGTTGTTGTATTAAATTTTCGGAC |
| CpG4MR1 | AATAAACTAAACCAAAAAAAACGAA |
| CpG4UF1 | TGTTGTTGTATTAAATTTTTGGATG |
| CpG4UR1 | AATAAACTAAACCAAAAAAAACAAA |
| CpG5MF1 | TTTCGTTTGTGTTTTTTTATTTGTC |
| CpG5MR1 | AAAATATACCTCAAAATCGATCGTT |
| CpG5UF1 | TTTGTTTGTGTTTTTTTATTTGTTG |
| CpG5UR1 | AAAATATACCTCAAAATCAATCATT |
